# A guide to trajectory inference and RNA velocity

**DOI:** 10.1101/2021.12.22.473434

**Authors:** Philipp Weiler, Koen Van den Berge, Kelly Street, Simone Tiberi

**Affiliations:** Institute of Computational Biology, Helmholtz Center Munich, Munich, Germany; Department of Mathematics, Technical University of Munich, Munich, Germany; Department of Applied Mathematics, Computer Science and Statistics, Ghent University, Ghent, Belgium; Bioinformatics Institute Ghent, Ghent University, Ghent, Belgium; Department of Statistics, University of California, Berkeley, CA, USA; Department of Data Science, Dana-Farber Cancer Institute, Boston, MA, USA; Department of Biostatistics, Harvard T.H. Chan School of Public Health, Boston, MA, USA; Department of Molecular Life Sciences, University of Zurich, Zurich, Switzerland; Swiss Institute of Bioinformatics, University of Zurich, Zurich, Switzerland

**Keywords:** Trajectory inference, RNA velocity, Single-cell RNA sequencing, Transcription, Splicing, Gene regulation, Cell differentiation, Bioinformatics, Computational biology

## Abstract

Technological developments have led to an explosion of high-throughput single cell data, which are revealing unprecedented perspectives on cell identity. Recently, significant attention has focused on investigating, from single-cell RNA-sequencing (scRNA-seq) data, cellular dynamic processes, such as cell differentiation, cell cycle and cell (de)activation. Trajectory inference methods estimate a trajectory, a collection of differentiation paths of a dynamic system, by ordering cells along the paths of such a dynamic process. While trajectory inference tools typically work with gene expression levels, common scRNA-seq protocols allow the identification and quantification of unspliced pre-mRNAs and mature spliced mRNAs, for each gene. By exploiting the abundance of unspliced and spliced mRNA, one can infer the RNA velocity of individual cells, i.e., the time derivative of the gene expression state of cells. Whereas traditional trajectory inference methods reconstruct cellular dynamics given a population of cells of varying maturity, RNA velocity relies on a dynamical model describing splicing dynamics. Here, we initially discuss conceptual and theoretical aspects of both approaches, then illustrate how they can be combined together, and finally present an example use-case on real data.

## 1. Introduction

In biology, we have been successfully organizing cells into different cell types, representing tissues which in turn constitute organisms. However, oftentimes the relationships between different cell states are not discrete, but continuous as in the case of cell differentiation or embryogenesis. Modelling these continuous spectra of cell states is the goal of trajectory inference (TI). Unlike clustering, which is the task of assigning categorical labels to cells, trajectory inference assigns continuous values, called pseudotime [1].

Pseudotime values are designed to quantify a cell’s position along a spectrum (Figure 1). In the case of differentiation, or any process in which progression occurs in a single direction, a cell’s pseudotime value can be viewed as a proxy for how far along the spectrum a cell is, relative to the progenitor state. However, pseudotime is based solely on transcriptional information, so it cannot be interpreted as an estimator of the true time since initial differentiation. Rather, it is a high-resolution estimate of cell state, which is likely to be monotonically related to the true chronological time, but there is no guarantee that equivalent changes in transcriptional profiles follow a similar chronological time.

**Figure 1.**
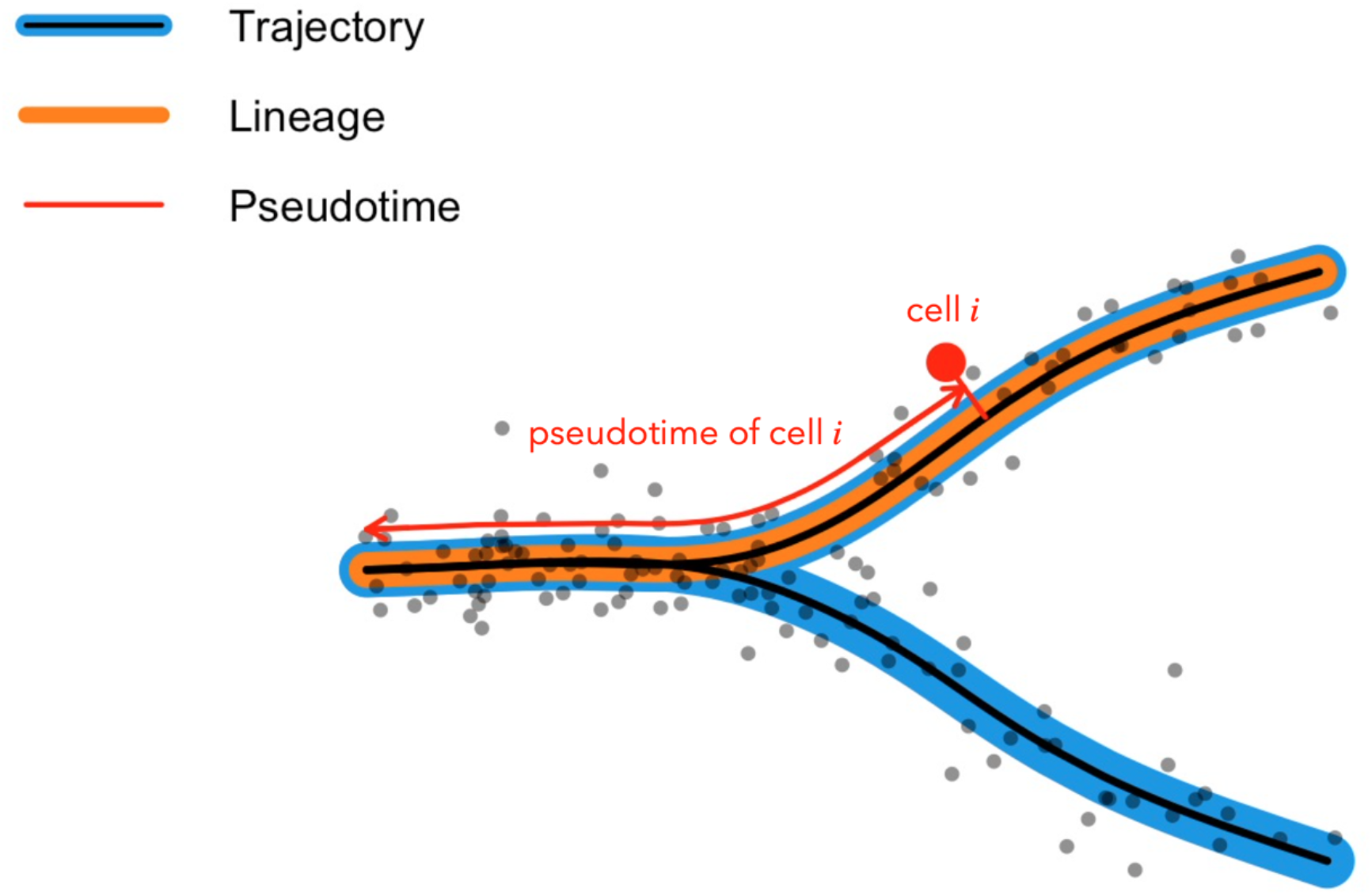
Introduction to trajectory inference terminology. A trajectory is a collection of (one or more) lineages. A lineage is a single differentiation path. The pseudotime of a cell is the length of the lineage from its inception up to the projection of the respective cell onto that lineage.

In Section 2 we will discuss the main conceptual ideas behind trajectory inference, including recommended upstream analyses. We will briefly introduce several types of trajectories, starting from a simple trajectory consisting of a single lineage (e.g., the differentiation of a unipotent stem cell to its mature cell state).

In the following, we will refer to a *trajectory* as a collection of lineages, which together describe a dynamic process (Figure 1). A cell’s pseudotime, then, is defined as the length of the lineage from its inception up to the projection of a cell onto that lineage. If a trajectory consists of multiple lineages, cells may have multiple pseudotime values, depending on which lineage the cell *is projected upon*.

Conversely to TI, RNA velocity describes the temporal change in mRNA over time by leveraging splicing dynamics [2]. These can be split into three main stages: transcription, splicing and degradation. A schematic overview of the process is given in Figure 2. During transcription, DNA is synthesized into precursor messenger RNA (pre-mRNA). Pre-mRNA contains both expressed regions (exons), as well as non-coding regions (introns) which are redundant for translation. Consequently, introns are removed through splicing. This leaves only exons forming mature mRNA. Finally, the process is completed with the eventual degradation of mRNA. In the following, we will refer to pre-mRNA as unspliced mRNA *u*, and to mRNA as spliced mRNA *s*. The rates of transcription, splicing and degradation are denoted by *α, β* and *γ*, respectively. Using this naming convention, the process of splicing has previously been modelled for each gene, independently of all others, by the following system of ordinary differential equations (ODEs) [3]

**Figure 2.**
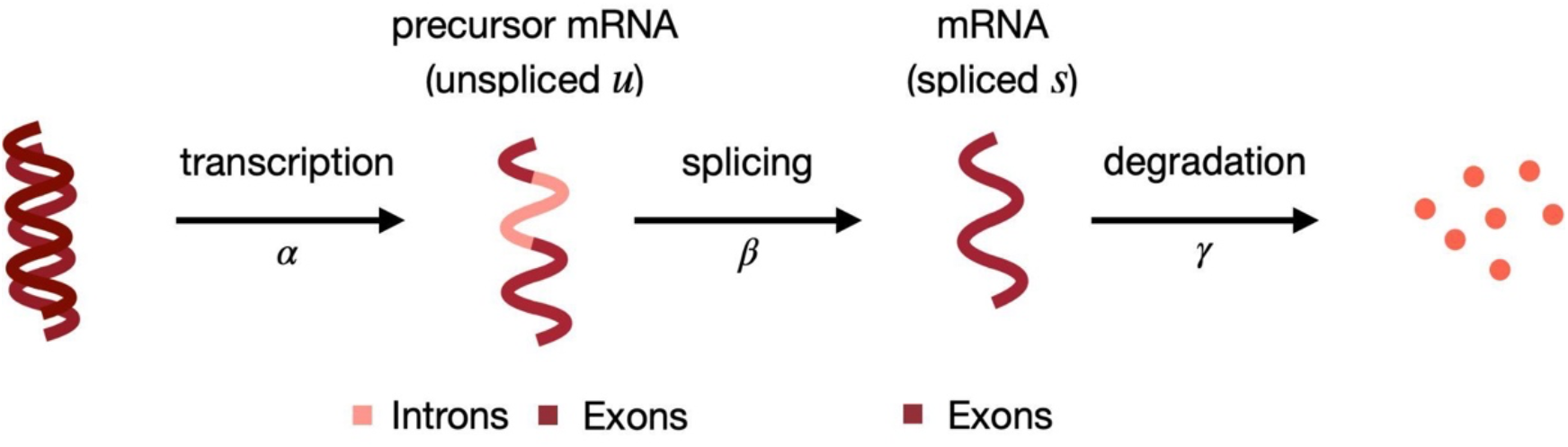
The first step in splicing dynamics consists in transcribing DNA to form unspliced, precursor mRNA, u. The included non-coding regions (introns) are removed through splicing. This process leaves only coding regions (exons) which form spliced mRNA, s, and are eventually degraded.

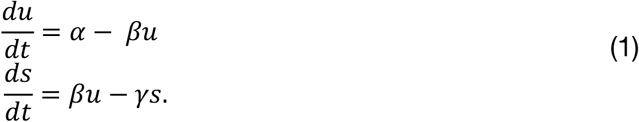

By definition, the RNA velocity is the derivative of spliced counts with respect to time.

The manuscript is organized as follows. Section 2 and 3 introduce the conceptual and theoretical aspects of TI and RNA velocity, respectively. Section 4 briefly explains how one can combine the two approaches by inferring trajectories from the RNA velocity output. Section 5 presents an example usage, on a real dataset, of both TI and RNA velocity. Section 6 reports some notes and practical guidelines for the actual implementation of both frameworks.

## 2. Trajectory Inference

### 2.1. Data pre-processing

Trajectory inference is most commonly applied to complex, large single-cell transcriptome sequencing datasets. Defining the trajectory in such high-dimensional datasets can be challenging, and therefore often calls for upstream analyses that allow us to uncover the major biological variation. We assume that we are working with a high-quality dataset, where each measurement refers to a single cell, i.e., where empty droplets, doublets/multiplets, as well as low-quality cells have been removed. First, we should account for major sources of technical variation, such as differences in sequencing depth between individual cells, using normalization. Often, cell-specific size factors are calculated to account for this [4]. Since each cell has thousands of dimensions, even on normalized data, it is difficult to visualize the global biological structure of the dataset without using dimensionality reduction techniques. Indeed, owing to the so-called *curse of dimensionality*, all cells tend to become approximately equidistant in high-dimensional space [5]. To alleviate this issue, we try to reduce dimensionality using feature selection, where we select a limited subset of genes which we believe still explain a substantial amount of variation in our dataset. Indeed, one may view feature selection as the most basic dimensionality reduction technique. Finally, in order to infer the trajectory, we attempt to summarize and visualize the dynamic process in a low-dimensional space using dimensionality reduction techniques, such as principal component analysis (PCA), uniform manifold approximation and projection (UMAP) [6] or t-distributed stochastic neighbour embedding (t-SNE) [7]. Care must be taken to properly take into account the (count) properties of single-cell datasets, in order to ensure that this low-dimensional space is driven by biological signal, and not technical variation as shown in [8,9].

### 2.2 Single lineage

Many biological systems can be adequately described by a single continuum, such as one cell type transitioning into another or the spatial distribution of related cells along an organ. We can model these systems with a trajectory structure consisting of a single lineage. In this case, the trajectory inference problem can be seen as a dimensionality reduction to a single dimension: we want to find a single dimension that captures as much biologically relevant variation as possible.

In this framework, cells’ loadings along the first principal component could be interpreted as pseudotimes. However, more specialized approaches generally provide better results. One reason for this is that the first principal component (PC1) may not always identify biological effects (e.g., sequencing depth or the fraction of zeros in each cell are often correlated with PC1) [9]. Similarly, PC1 captures distinct proportions of the overall variance in different datasets. Another issue is the so-called “horseshoe effect,”, where a continuous gradient appears as a U-shaped curve in the first two principal components [14].

Beyond PCA, many trajectory inference methods exist and are sufficiently adaptive to appropriately model a single lineage [15]. However, few are able to accurately infer directionality. With one lineage, this is a binary decision: which of the two ends should be the starting point? Without bringing in additional data or domain-specific knowledge, this can be a difficult question to answer, hence most trajectory inference methods rely on users’ supervision. A more data-driven approach may require annotated datasets and/or cell-type identification methods to select the cluster that most closely resembles stem cells; this cluster is then used as the starting point of the trajectory.

### 2.3 Diverging lineages

Since trajectory inference is often applied to populations of differentiating cells, we are frequently interested in modeling multiple branching lineages. In this case, we not only need to determine each cell’s pseudotime, but also its degree of commitment to a particular fate. Consider a bifurcating trajectory (i.e., consisting of two diverging lineages); cells in the starting population have not yet committed to a fate, whereas cells in the terminal populations have. In between these two extremes, there are likely to be different degrees of commitment to each of the two possible endpoints.

By their nature, these sorts of trajectories lend themselves well to tree-like structures. Hence, many TI methods construct a branching graph to summarize the layout of major cell types, with the root node representing the initial, undifferentiated population and (other) leaf nodes representing terminal populations [16–18]. For example, on the pancreas dataset [19] employed in Section 5, we used *Slingshot* [17] to construct a supervised graph on the set of cell clusters, resulting in a trajectory with four possible fates.

### 2.4 Converging lineages

In some systems, cells from different progenitor populations converge into a single cell type. Any TI method that can model diverging lineages can also model simple converging lineages, by selecting the terminal population as the “beginning” of the trajectory and then reversing the resulting pseudotimes. However, more complex systems that involve both divergence and convergence can be difficult to model and require methods that accommodate such complexity, such as *PAGA [20]* and *Monocle3* [18].

### 2.5 Cycles

Another prominent feature of some trajectories is the cell cycle. On its own, a cyclic trajectory can be modelled in many ways, but as part of a larger trajectory, these can be difficult to accurately represent. Not only do cyclic components break the tree-like graph structure assumed by many TI methods, but they also introduce some ambiguity regarding the interpretation of pseudotime. Specifically, a cell which has undergone one full cycle may be transcriptionally indistinguishable from a cell which has not undergone any changes; this may invalidate the interpretation of pseudotime as being monotonically related to real time.

For instance, in the pancreas dataset (Section 5), there is a noticeable cycle in the first two clusters. However, since *Slingshot* assumes a tree-like structure, it is unable to accurately model this part of the trajectory.

### 2.6 Challenges and limitations

All trajectory inference procedures attempt to recover a ‘best-fitting’ trajectory on any dataset to which they are applied, just like *k*-means clustering always provides *k* clusters. Care must be taken that the structure one attempts to model indeed represents a biological dynamic process. Poor preprocessing or improperly accounting for technical variation may give rise to spurious trajectories; for instance, doublet cells may be misinterpreted as ‘intermediate states’ [21], or a hypothesized major source of biological variation may actually be reflecting technical effects, such as the fraction of zeros or the total number of counts for each cell [9].

For some datasets, there is a thin line between clustering the cells into discrete entities and modeling the structure as a trajectory. For example, depending on the parameters of one’s preferred dimensionality reduction technique, the structure may seem discrete rather than continuous. Therefore, before estimating a trajectory on any dataset, domain-specific knowledge remains irreplaceable, e.g., through close examination of well-characterized marker genes. While computational algorithms may indeed be able to fit the observed structure, they will not be able to determine whether or not it is biologically plausible, and results may be misleading.

## 3. RNA velocity

### 3.1 Data pre-processing

RNA velocity analyses are based on the counts of unspliced and spliced mRNA. Pipelines such as velocyto [2], alevin [10], alevin-fry [11] or kalisto/bustools [12] allow inferring them from scRNA-seq data. As in classical scRNA-seq analysis, measurements need to be pre-processed [13]. First, genes are filtered by expression level or by their occurrence in cells. Only genes with enough counts in both modalities should be retained. Following this first filter, each modality is normalized, and highly variable genes are identified and extracted, while all remaining genes are discarded.

Single-cell RNA sequencing data is notoriously noisy. To extract the relevant signal and infer RNA velocity, it needs to be smoothed first. This is done by calculating the first order moment (mean) of an observation over its k nearest neighbours. Figure 3 shows how the phase portrait of gene *Cpe* changes, in a pancreatic endocrinogenesis dataset [19] (see Section 5), with an increasing number of neighbours. Whereas there is no curvature present in the raw data (leftmost plot), the up-regulation in gene *Cpe* becomes visible when increasing the number of neighbours. This illustrates the importance of data imputation by moments for RNA velocity analysis. The phase portrait not changing dramatically after a certain number of neighbours *k* indicates robustness with respect to the choice of this parameter. Increasing *k* to large values is not recommended as it increases computational complexity and does not describe well the most similar observations. For large values of *k*, observations not similar to the reference would, thus, be included as well.

**Figure 3.**
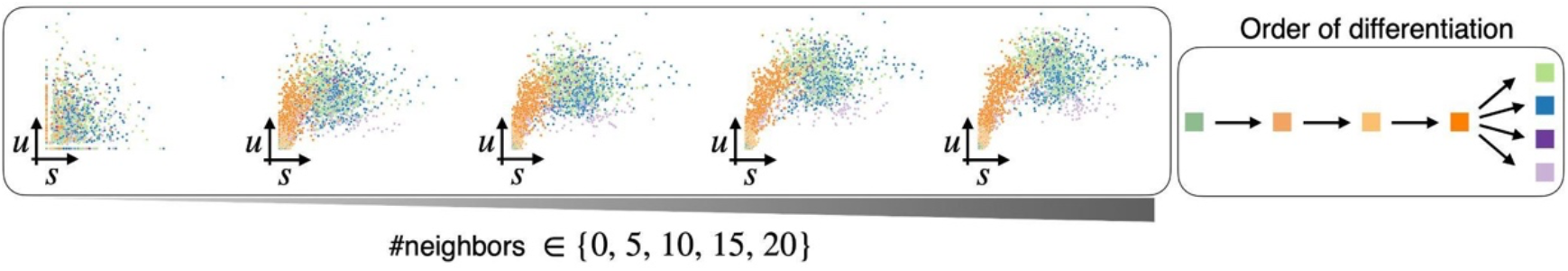
Imputing measurements by first order moments removes noise. With an increasing number of neighbours (from left to right), the phase portrait becomes less noisy and up-regulation is visible. Starting from the origin, time increases when moving along the trajectory clockwise. Consequently, cell types should be ordered along the trajectory accordingly. Comparing this ordering of cell types with the biologically known order of differentiation, an up-regulation is identified as the correct regulation type.

### 3.2 Splicing dynamics

The solution of the simple, linear system (1) can be found analytically and is given by

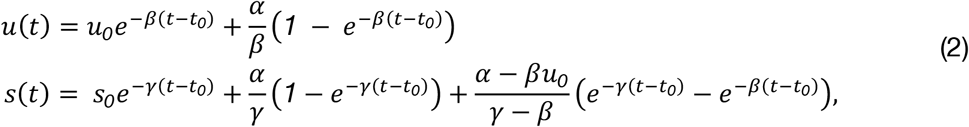

when *γ* ≠*β*, and where *s*_*0*_ and *u*_*0*_ refer to the unspliced and spliced counts at time *t*_*0*_, respectively, and *e*. indicates the exponential function.

Understanding how splicing dynamics evolve over time is fundamental for analyzing RNA velocity. During transcription, each modality increases until it reaches a steady state. As spliced mRNA is produced from unspliced mRNA, the abundance of the former increases prior to the latter. If transcription ends prematurely, steady states may not be reached. Using the proposed system of ODEs (2), this process can be simulated and abundances plotted against time (Figure 4a). However, this cannot be done in real data measurements due to the destructive nature of scRNA-seq experiments. Consequently, phase portraits (Figure 4b), where the abundance of unspliced mRNA is plotted against the one of spliced mRNA, are studied, instead. In the ideal case, the full process of transcription, splicing and degradation is observed leading to an almond shaped phase portrait. The upper arc of the almond shape corresponds to the induction phase ending in a steady state in the upper right part of the portrait. Conversely, cells in the lower arc are in repression, i.e., transcription is turned off. If only part of the full cycle is observed during the experiment, the resulting portrait contains only the corresponding portion of the almond shape. Overall, this shape depends on the model parameters.

**Figure 4.**
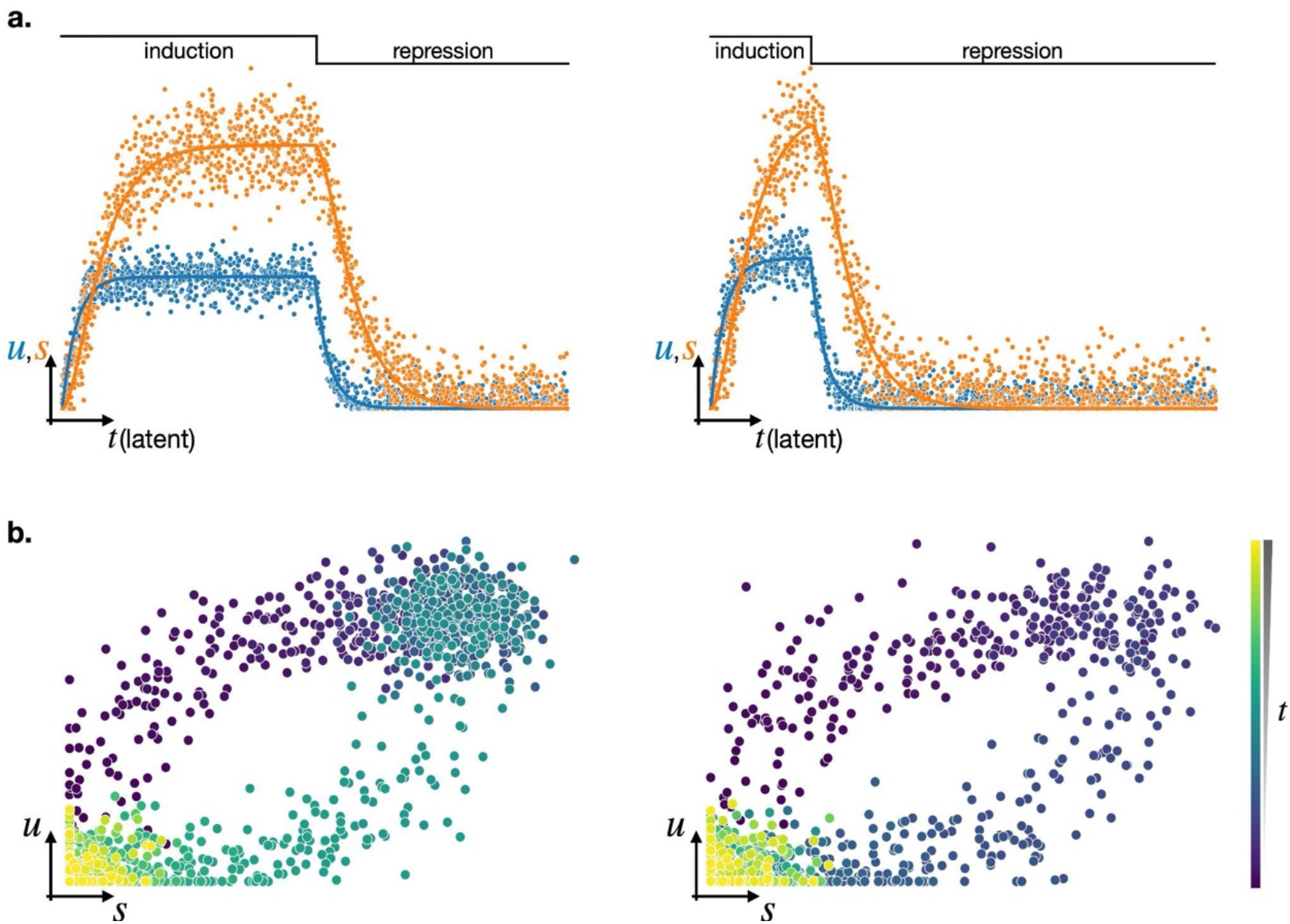
Visualization of measured abundances. **a**. Since spliced mRNA s is produced from unspliced mRNA u, the abundance of s increases (or decreases) with a temporal delay with respect to u. During DNA transcription (i.e., induction phase), both s and u increase. If this phase lasts long enough, a steady state is reached (left). Then, during repression, transcription is turned off and both modalities gradually decrease. **b**. As single-cell measurements are snapshot data, time can neither be tracked over several measurements nor be associated to a single one. Unspliced and spliced counts are plotted against each other in a phase portrait. Its curvature increases as the temporal delay between s and u grows.

In the following, we will first discuss theoretical aspects of the RNA velocity analysis. This includes how scRNA-seq should be processed, two methods for inferring model parameters and finally some RNA velocity based downstream analyses. Then, in Section 5.2, these discussed facettes will be applied to and showcased in a real-world dataset of endocrine development in the pancreas [19].

### 3.3 Parameter inference

Rate parameters cannot be measured during a scRNA-seq experiment and need to be inferred instead. Since scRNA-seq data consist of snapshot measurements, time can neither be measured nor tracked. Consequently, time points assigned to each cell, as well as the time when the system switches from induction to repression, need to be estimated.

This Section discusses how parameters are inferred and is divided into two parts: the first one discusses the estimation of the steady-state ratio using the so-called steady state model, the second one outlines how the full set of observations and model parameters can be inferred using a different approach [2,22].

#### 3.3.1 Steady-state model

In a first attempt to estimate RNA velocity, splicing dynamics are assumed to have reached their steady states [2]. In these steady states, we consequently find that:

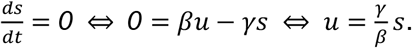

Note that this equation is only satisfied for cells in equilibrium, i.e., cells found in the lower and upper quantiles of the phase portrait. Assuming a common unit splicing rate *β* across all genes, we can estimate the steady state ratio, 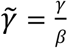, through an extreme-quantile linear regression with closed form solution

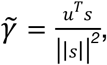

Where ∣| · |∣ indicates the euclidean distance. The residual to the linear-regression fit of the unspliced mRNA in cell *c* (Figure 5a), that we denote by *v*_*c*_, is defined as

**Figure 5.**
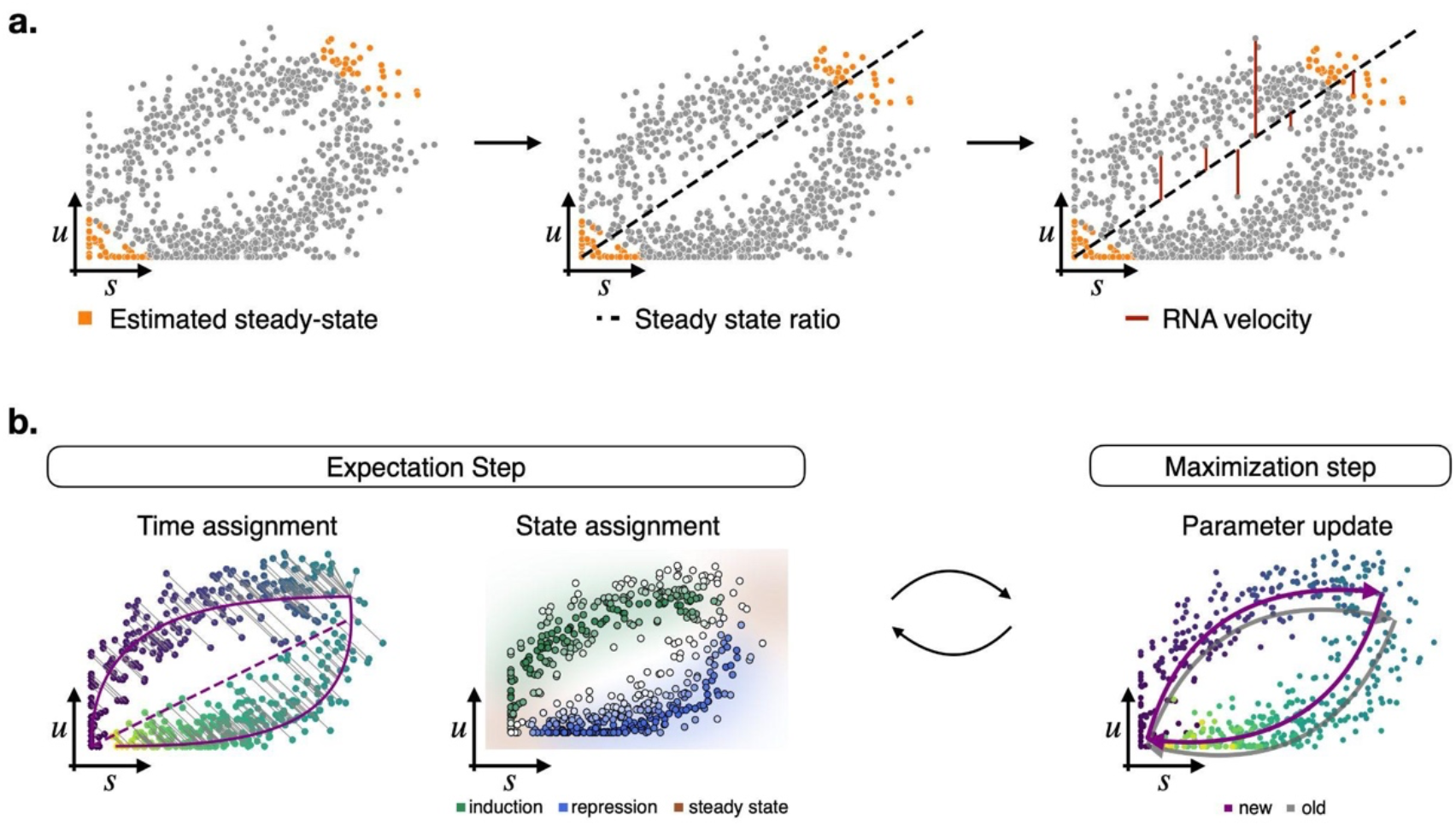
Inferring RNA velocity and model parameters from scRNA-seq data. **a**. The steady state model assumes the system has reached its steady states and estimates the cells that have reached either steady state (left). A linear regression on these extreme quantiles is fitted (middle), and the RNA velocity is estimated as the residuals to this fit (right). **b**. Solving the full splicing dynamics allows generalizing to transient genes. Parameters are inferred by expectation-maximization (EM). During the E-step (left), each observation is assigned both a time and a state (induction, repression, or steady state). The former assignment is found by minimizing the distance to the current estimate of the trajectory, the latter by the maximal likelihood of a state. Then, model parameters are updated by maximizing the overall likelihood during the M-step (right).

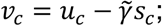

this residual is the estimated RNA velocity of cell *c*.

In other words, this simple estimation of RNA velocity requires both steady states of the system to be observed, as well as a common splicing rate *β* across all genes. While this approach proves fruitful in some cases, its assumptions represent serious limitations for its applicability. On one hand, the estimated steady state ratio will deviate from the real (unobserved) ratio when not all steady states are observed; this occurs, for example, when transcription ends prematurely. On the other hand, splicing is a gene-dependent property, and its rates can largely vary between genes. Furthermore, not inferring the full set of model parameters and only using a subset of the available data are additional drawbacks. Below, we discuss how these shortcomings can be addressed and resolved.

#### 3.3.2 Dynamical model

Having preprocessed the scRNA-seq measurements, we are interested in inferring model parameters *α, β, γ* of (1). Since scRNA-seq experiments are destructive by nature (i.e., cells are destroyed when sequenced, and therefore cannot be measured multiple times), observations cannot be associated to a specific time point. As such, the observations contain latent, i.e., hidden or unknown, variables; these are time point *t* and state *k* (induction, repression, or steady state) of an observation. Consequently, we cannot estimate model parameters via maximum likelihood, but turn to expectation-maximization (EM) [23], as proposed by [22] and implemented in the Python package *scVelo*, instead. The algorithm is divided into two steps - the E-step (i.e., expectation step) and the M-step (i.e., maximization step). Hidden variables are updated during the E-step, while the remaining parameters are updated during the M-step (Figure 5b). To infer model parameters, we assume constant rates across time. This requires splitting (2) into two parts: induction and repression. During induction the transcription rate *α* is positive, while during repression it is equal to zero. Finding the optimal time assignment is computationally expensive. To boost performance, the optimal solution can be approximated, leading to a 30-fold speed-up [22]. For this, we rewrite the analytic solution (2) for *s* as

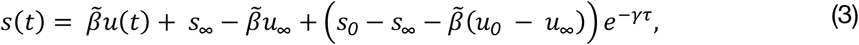

with 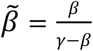, and *s*_∞_ and *u*_∞_ being the values of the systems at equilibrium (i.e., as time goes to infinity). Defining 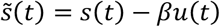 and initial and steady states accordingly, (3) can be solved for *τ* as

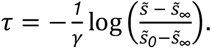

In case *β*≤ *γ*, the inverse of *u*(*t*) is considered instead, i.e.,

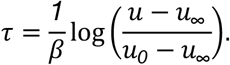

The EM algorithm estimates gene-specific reaction rates and assigns gene-specific time points to each observation. These parameters are only identifiable up to a proportionality constant *λ*. It is easy to see that multiplying all rates by *λ* and dividing time by the same value does not change the solution. If the true time scale [*t*_*0*_, *t*_*max*_] was know, the scaling parameter *λ* could be factored out. Here, *λ* is estimated by aligning the inferred time assignment across all genes to a common scale.

### 3.4 From velocities to transition probabilities

RNA velocities, representing the true underlying splicing dynamics, allow predicting the future state of cells. Given the gene profile of cell *j* at time *t, c*_*j*_, and its future gene profile at time *t* + *Δt* (with *Δt* > *0*), 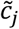, the change in its gene profile is estimated by 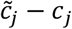. Since future gene profiles cannot be assigned to each cell, their neighbours are considered as candidates. The probability of transitioning to one neighbour is high if the heuristic displacement and velocity vector point into a similar direction (Figure 6a). This similarity is captured by cosine similarity, i.e.,

**Figure 6.**
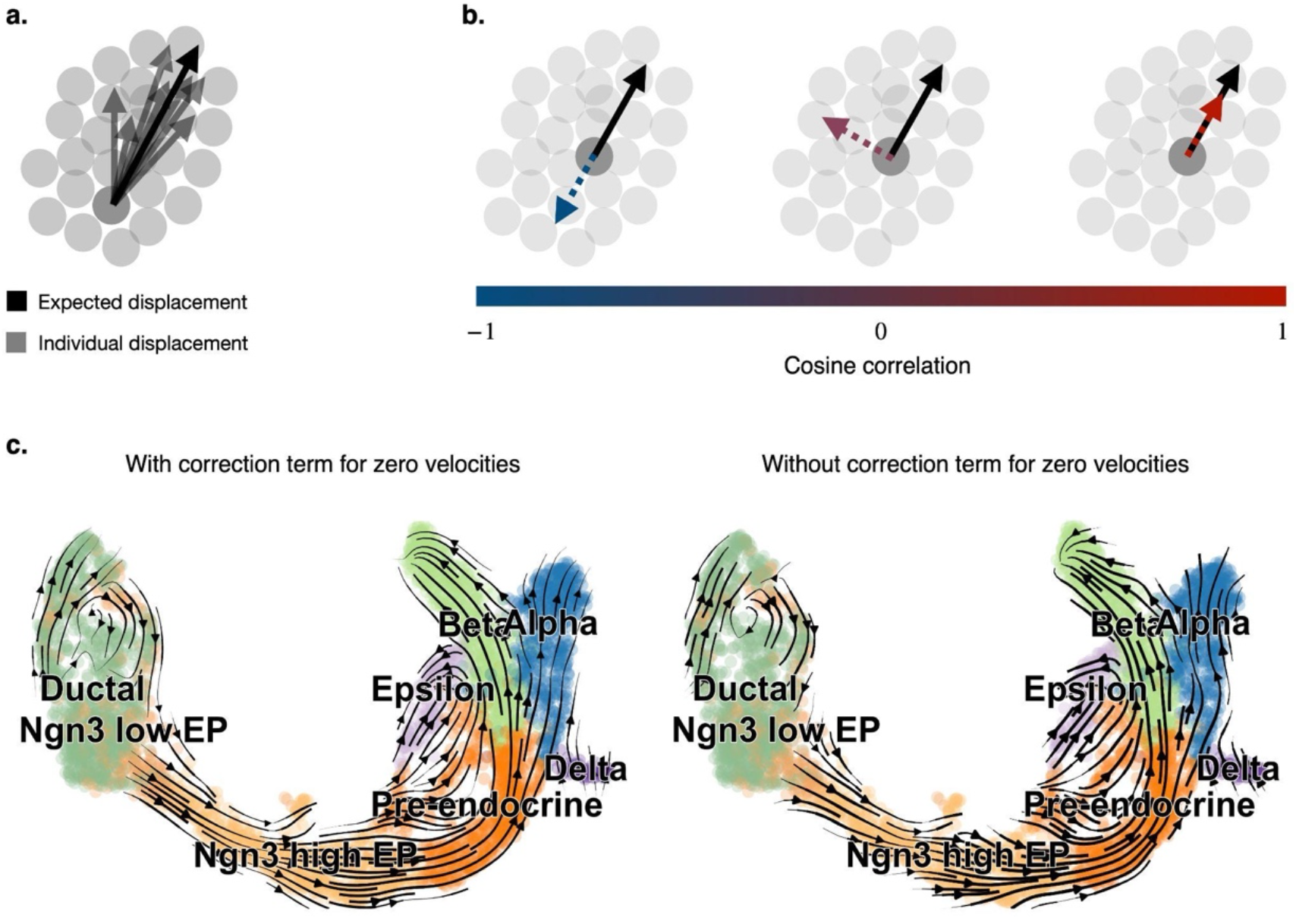
Visualization of velocities (application example taken from [22]). **a.** The expected displacement of a reference cell indicates the direction of its future state. Here, the expected displacement is given by a weighted average of the displacement between a reference and its neighbours. Transition probabilities are used as weights. **b**. The cosine correlation measures the similarity between two vectors. Here, displacement and velocity vectors are compared to estimate transition probabilities. **c**. To account for vanishing or orthogonal velocities, the expected displacement is corrected by the mean displacement (left). The correction term particularly affects, compared to omitting it (right), the border of the embedding.

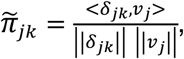

with displacement *δ*_*jk*_ = *c*_*k*_ − *c*_*j*_ and velocity vector *v*_*j*_ for cell *j*. As visualized in Figure 6b, 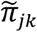 is maximal when displacement and velocity vector point into the same direction. In case of orthogonality or opposite directions, 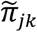 vanishes or takes its minimum, respectively. The graph associated with this similarity matrix is called velocity graph.

Similarity measures cannot necessarily be interpreted as probabilities. The cosine similarity, for example, falls in the interval [−*1, 1*]. An exponential kernel is applied to transform cosine similarities to transition probabilities, i.e.,

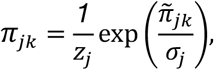

with normalization constant *z*_*j*_ and parameter *σ*_*j*_ adjusting the kernel width. These transition probabilities define the elements of the transition matrix *Π*, which allows determining descendants and ancestors of a given cellular distribution. Visualizing these in a low dimensional embedding of a latent space can be a powerful tool to easily and intuitively study cellular differentiation. Common latent space representations are UMAP and t-SNE [6,7]. However, two- or three-dimensional representations should be interpreted with caution as they contain limited information compared to higher dimensional representations [24]. As such, they should not form the sole basis of quantitative modeling.

Differentiation paths can be visualized by an adjusted expected lower-dimensional displacement. Given the embedding vectors *s*_*j*_and *s*_*k*_, the expected displacement is given by

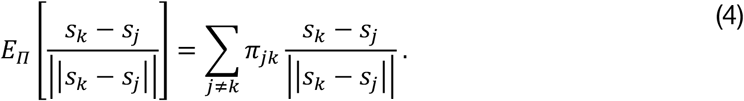

In case of vanishing or orthogonal (with respect to the displacement) velocities, transition probabilities are uniform and the heuristically expected displacement is zero; nonetheless, the value of (4) is not null. For consistency, (4) is therefore adjusted as:

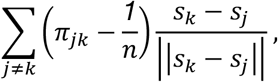

with *n* being the number of possible transitions (Figure 6c).

### 3.5 Gene-shared latent time

The inferred model parameters and time assignments are gene specific. In reality, however, each cell is associated with a time. As such, gene-specific times need to be pooled together into a single, gene-shared latent time for every cell. To mitigate the effect of noise, only a subset of genes, for example well-fitted ones, are considered. In addition to the goodness of fit, the correlation between the inferred time and a velocity graph based pseudotime is a decisive factor for selecting genes.

To infer a velocity based pseudotime, as a first step, initial and terminal states are identified. They can heuristically be estimated by the stationary distribution *µ*\* satisfying

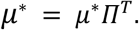

For a more sophisticated approach of identifying these states, the Python library *CellRank* can, for example, be used [25] (see Section 4).

Given an initial state, the average number of steps (in a random walk) to reach a given cell from the starting cell defines, similar to diffusion pseudotime [26], a pseudotime. Compared to this approach, diffusion pseudotime uses a similarity-based diffusion kernel and does not implicitly infer terminal states. Then, genes for which the correlation between their time assignments and the constructed pseudotime does not exceed a given threshold, are neglected from further analysis.

Finally, for each estimated terminal state, p-quantiles of time increments (with respect to a given initial state) across all genes are calculated. This is motivated by cell density being not uniformly distributed across time. Instead, the density usually increases in late time points. The gene-shared latent time is then given by the mean p-quantile across all starting cells.

### 3.6 Challenges and limitations

While RNA velocity has had tremendous success and has been applied successfully to various datasets, the current framework fails in some scenarios. Even though extending the steady-state model to solving the full splicing kinetics generalized the concept to transient populations, the model still relies on constant parameters. Examples of violations of this assumption are transcriptional bursts, i.e., when the transcription rate increases abruptly after some time, or multiple kinetics. Transcriptional bursts can, for example, occur in hematopoiesis [27,28], and multiple kinetics in dentate gyrus [2,22,29].

Another limitation of the current formulation is that it relies on phase portraits exhibiting at least a partial almond shape. In reality, gene measurements are noisy, i.e., do not always lead to such shapes in the phase portraits. In this case, model parameters may be inferred incorrectly leading to wrong conclusions. This can, for example, be shown on a dataset of peripheral blood mononuclear cells including only mature cells [29,30]. In this case, albeit no differentiation is expected, the velocity stream has a clear, but arbitrary, direction.

See [29] for a more in-depth discussion on existing pitfalls and possible extensions to the existing framework.

## 4. Combining trajectory inference and RNA velocity

Whereas RNA velocity assigns directed, dynamic information to each cell, trajectory inference models the continuous differentiation of cell types on the phenotypic manifold. To combine the two approaches, *CellRank* has been proposed for RNA velocity informed TI [25]. For this, *CellRank* describes state transitions as a Markov chain to, first, identify initial, intermediate and terminal states and, then, determine fate probabilities of each cell, as well as expression trends and putative decision drivers in different trajectories. Markov chain models consist of states and transition probabilities between states. In *CellRank*, each state corresponds to a single cell. The possible transitions and initial estimates of their probabilities are given by a similarity based k-nearest neighbour (kNN) graph. To not only rely on transcriptional similarity, *CellRank* considers the directional information provided by RNA velocity as well. For this, a transition matrix similar to the one defined in Section 3.4 is used and uncertainties are propagated. Instead of cosine correlation, *CellRank* centres gene expression and velocities to zero mean, resulting in Pearson’s correlation. Both transition matrices (i.e., the similarity and velocity based ones) are combined by a weighted mean for a more robust estimate of transition probabilities. While *CellRank* has originally been developed to combine RNA velocity with transcriptomic similarity, it has meanwhile been extended to include other estimates of directionality, such as pseudotime, as well.

Since scRNA-seq is high dimensional and noisy, the transition matrix cannot be used as defined to solve the problem of trajectory inference. Instead, the system is first coarse grained into states, so-called macrostates, forming initial, intermediate and terminal states. These states can be automatically classified on the basis of coarse-grained transition probabilities and the probability to transition into each terminal state, the fate probability, can be quantified as well. Finally, *CellRank* uses these results to study fate commitment or identify putative driver genes specific to individual trajectories.

## 5. Hands-on: real data analyses

### 5.1 Trajectory inference analysis pipeline

We demonstrate some of the ideas presented above with a hands-on analysis of a single-cell pancreatic endocrinogenesis dataset taken from [19]. We perform this analysis in R, making use of multiple Bioconductor packages, most notably *Slingshot* for the task of trajectory inference.

The data can be easily downloaded in a usable format here: https://github.com/theislab/scvelo_notebooks/tree/master/data/Pancreas/endocrinogenesis_day15.h5ad. After downloading the data, we read it into R as a SingleCellExperiment object via the *zellkonverter* package [31]. This creates an object that contains not only the count matrix, but also cluster labels and dimensionality reductions from the original publication.

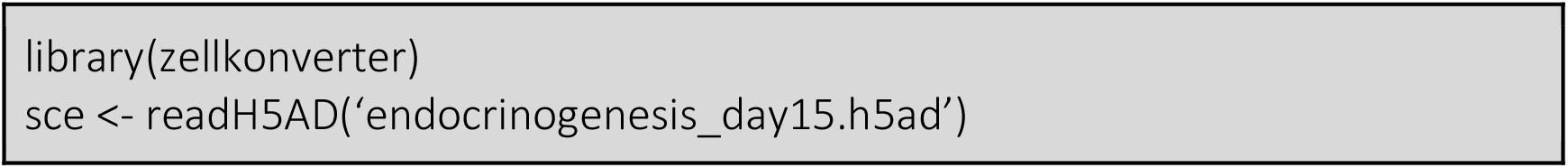

We are now ready to perform trajectory inference. We provide *Slingshot* with the UMAP dimensionality reduction and the cluster labels, as well as some biological supervision. In this case, the only supervision that impacts our results is the specification of the “Alpha” cluster as an endpoint; all other endpoints and the starting cluster are correctly inferred by *Slingshot*. We also increase the flexibility of the resulting curves by adjusting the “*spar*” parameter. This parameter is passed to the smoothing spline function which underlies *Slingshot*’s curves, and it allows us to account for the elongated nature of the UMAP, where most of the differentiation between lineages occurs toward the ends.

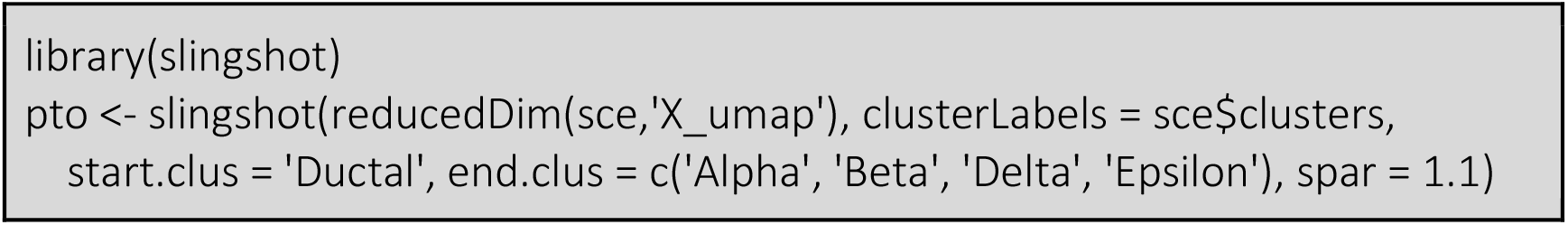

We can then plot *Slingshot*’s minimum spanning tree (MST) on the 2-dimensional UMAP (Figure 7).

**Figure 7.**
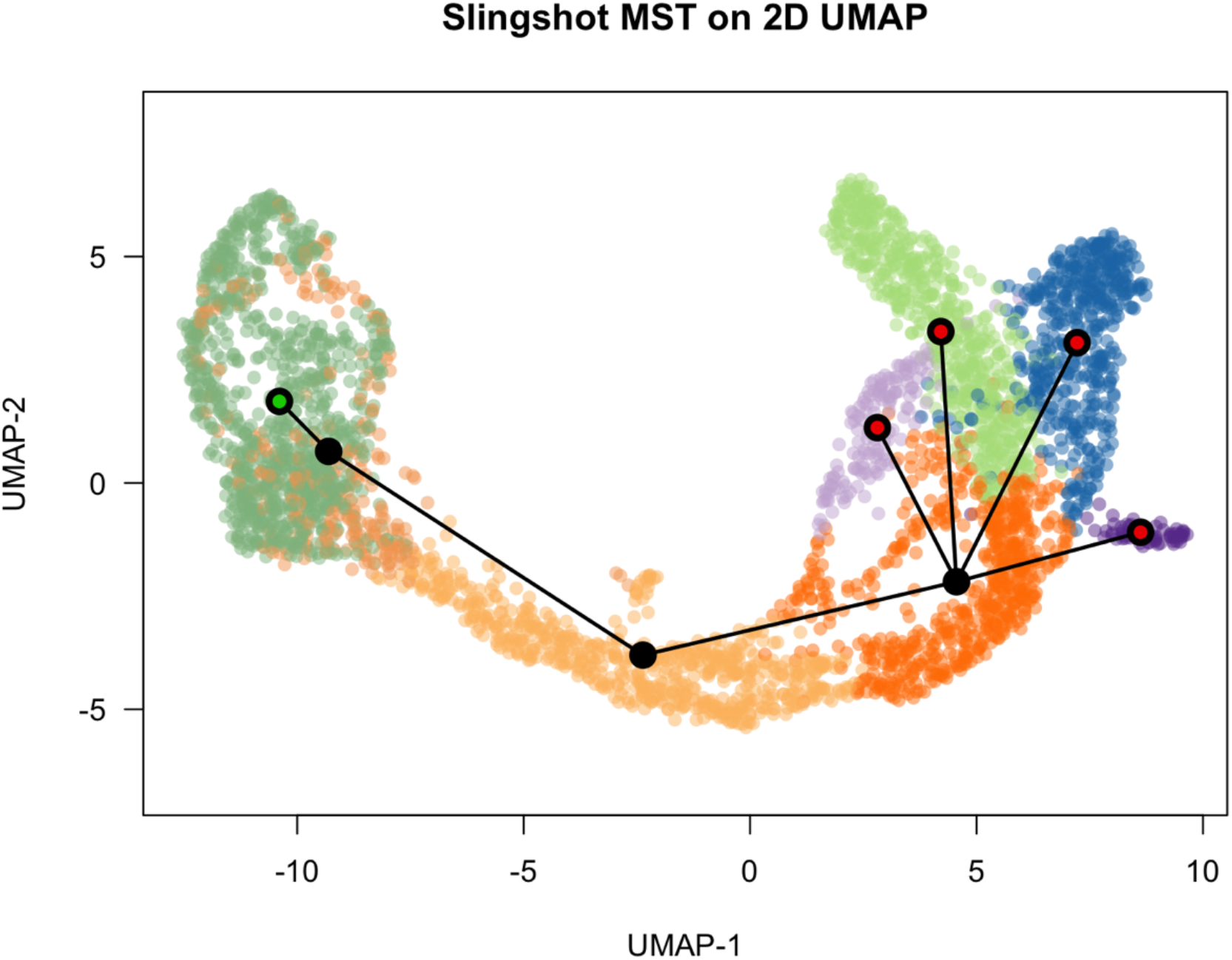
The minimum spanning tree, computed by Slingshot, visualized on a 2-dimensional UMAP embedding. Green (left), black (middle), and red (right) circles indicate initial, intermediate, and terminal clusters, respectively.

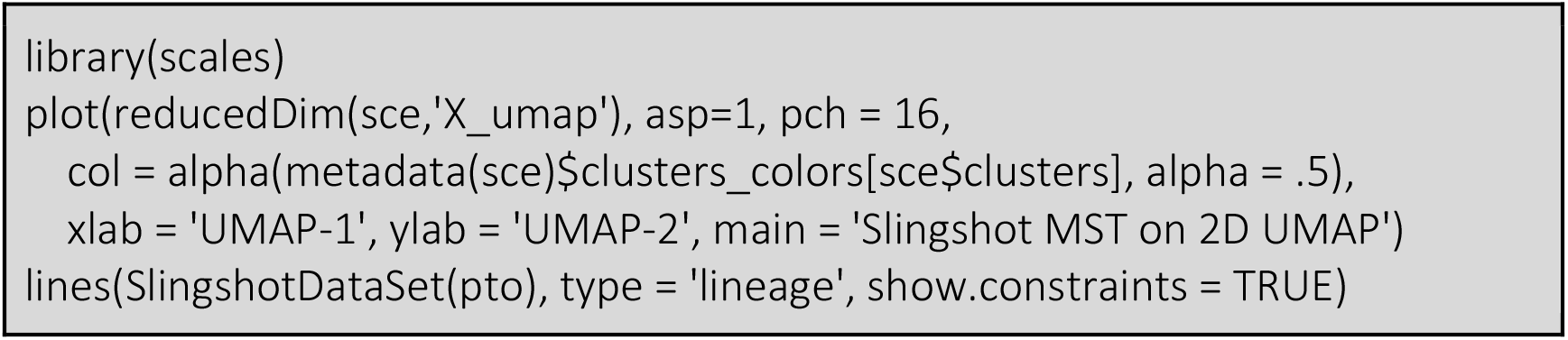

From the two-dimensional UMAP in Figure 7, we can see the cluster labels (representing cell types) and four distinct endpoints. However, there is significant overlap between the terminal clusters on the right-hand side of the plot, indicating that two dimensions might not be enough to fully characterize this complex trajectory structure. Therefore, we produced a three-dimensional UMAP (calculated from the pre-computed top 50 PCA components), and re-run *Slingshot* on this embedding. This provides us with a reduced-dimensional space in which the four terminal clusters are more clearly separated (Figure 8).

**Figure 8.**
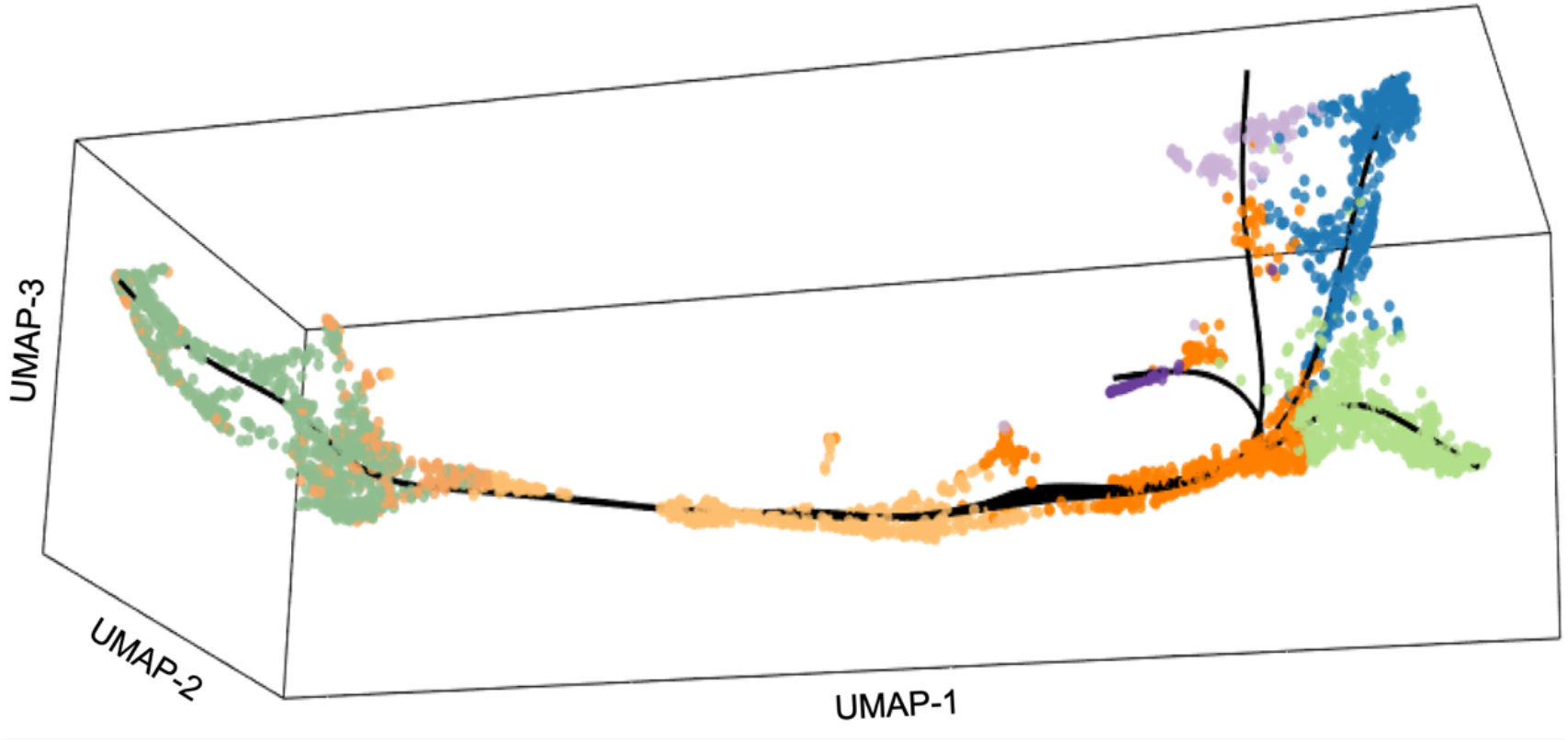
Visualization of the trajectory computed by Slingshot, on a 3-dimensional UMAP embedding.

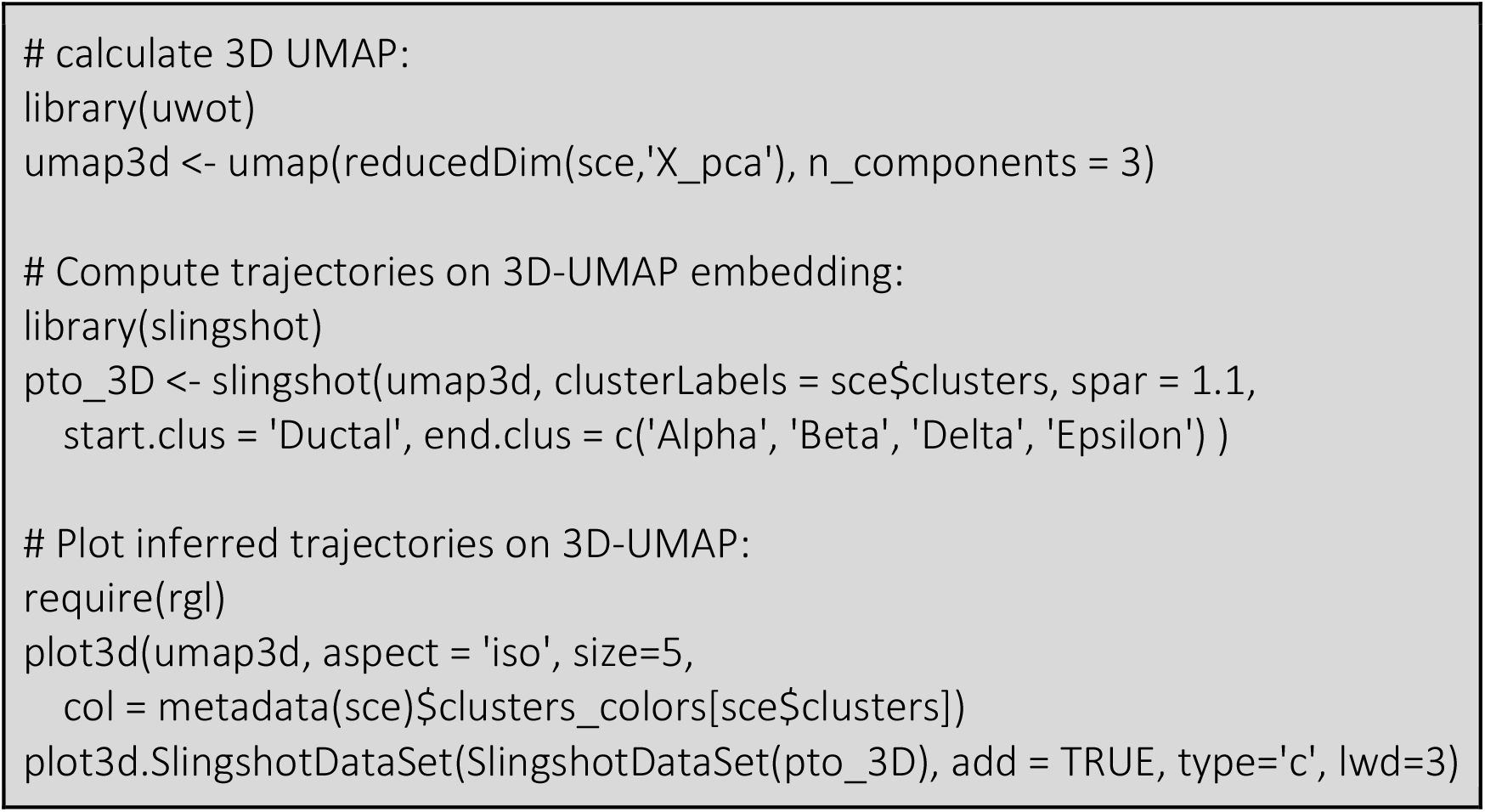

For visualization purposes, we also produce an embedding of these curves in the original, two-dimensional space. Note that this process does not change the pseudotime values, but is merely a way to visualize our results in a different space. The sharp turn in the Delta lineage (Figure 9) reinforces our earlier notion that the two-dimensional UMAP embedding was insufficient to adequately represent this trajectory.

**Figure 9.**
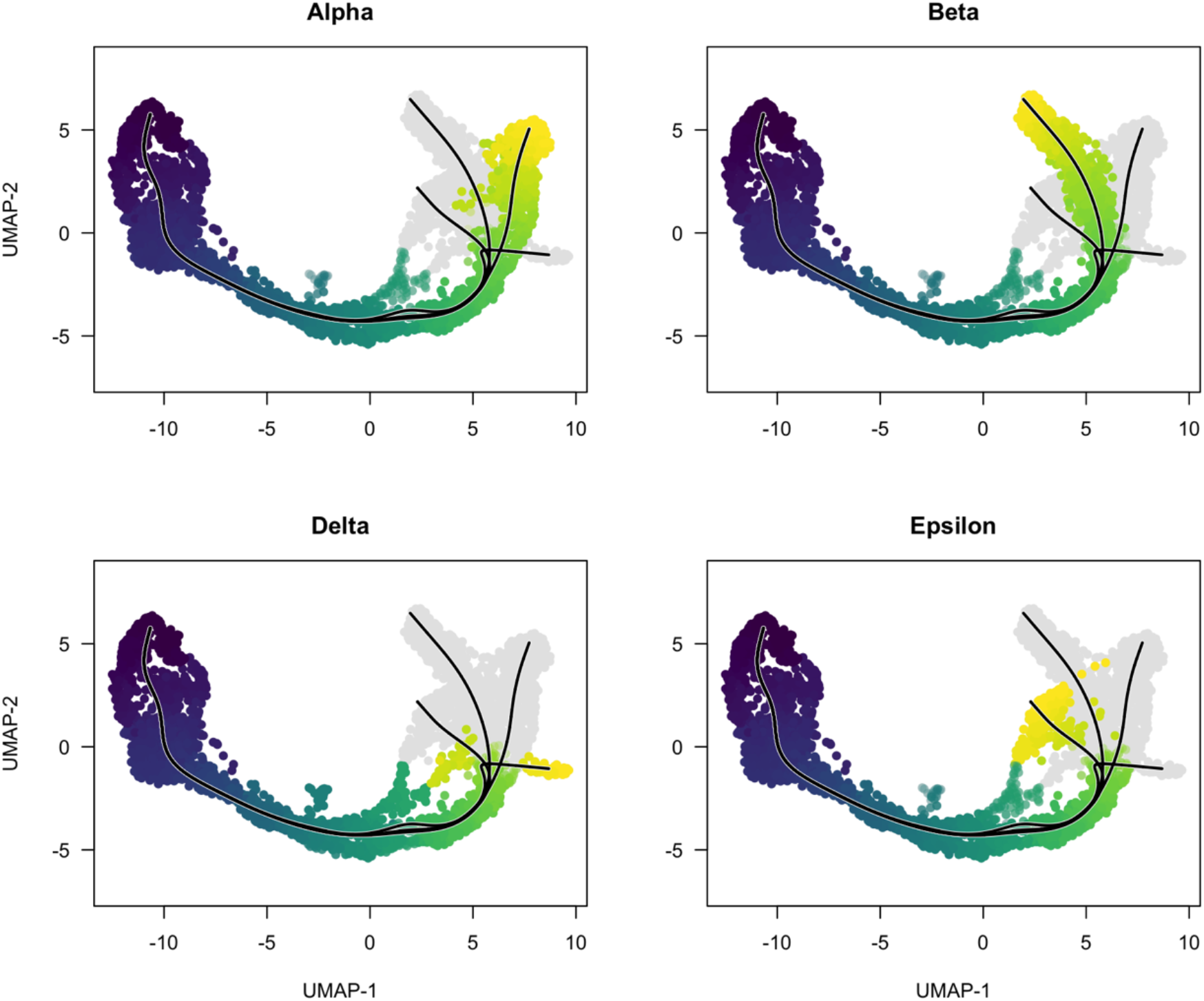
The trajectory and pseudotime computed by Slingshot, visualized on a 2-dimensional UMAP embedding. Each image refers to an end cluster: Alpha (top left), Beta (top right), Delta (bottom left), and Epsilon (bottom right).

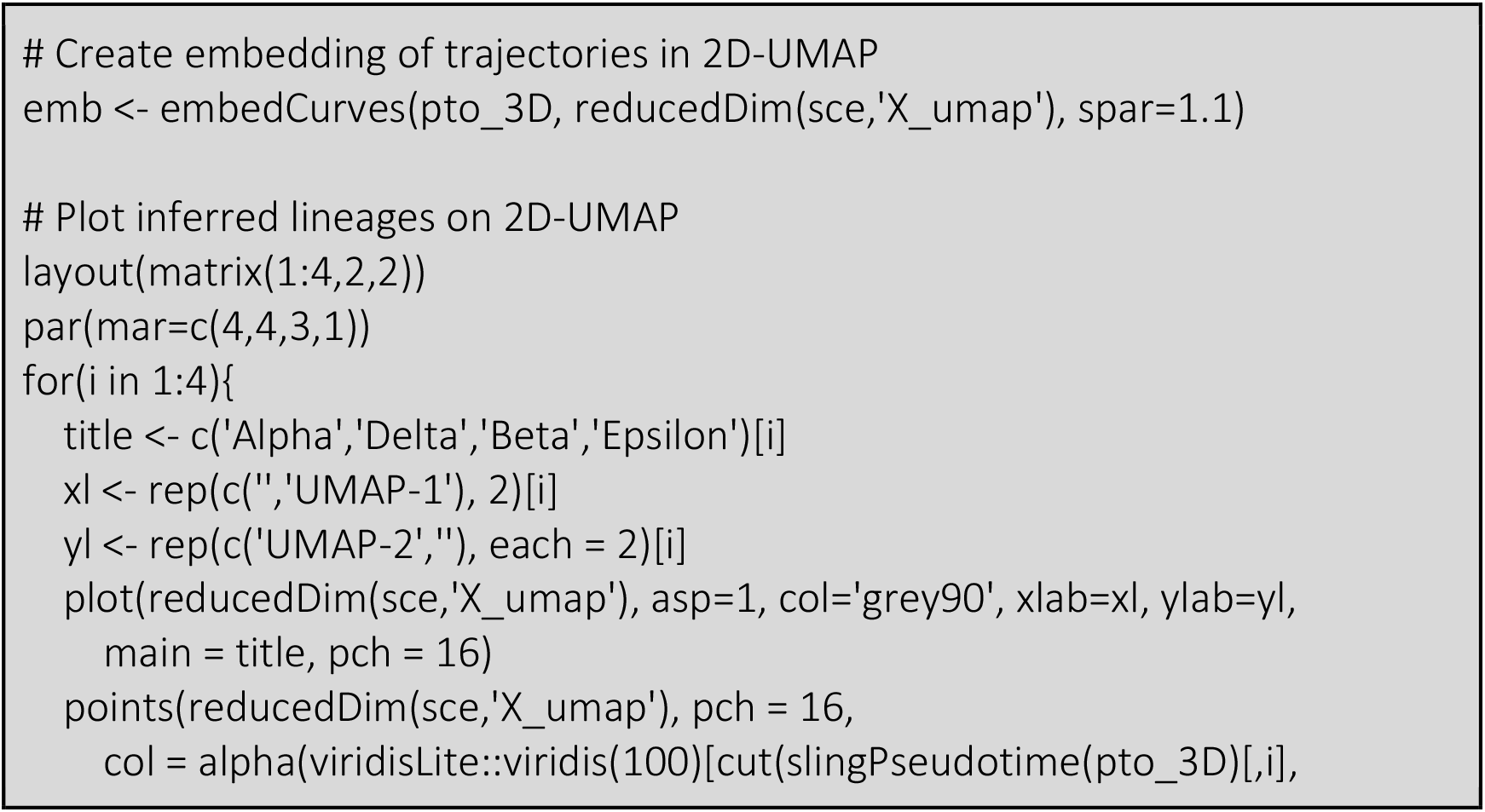
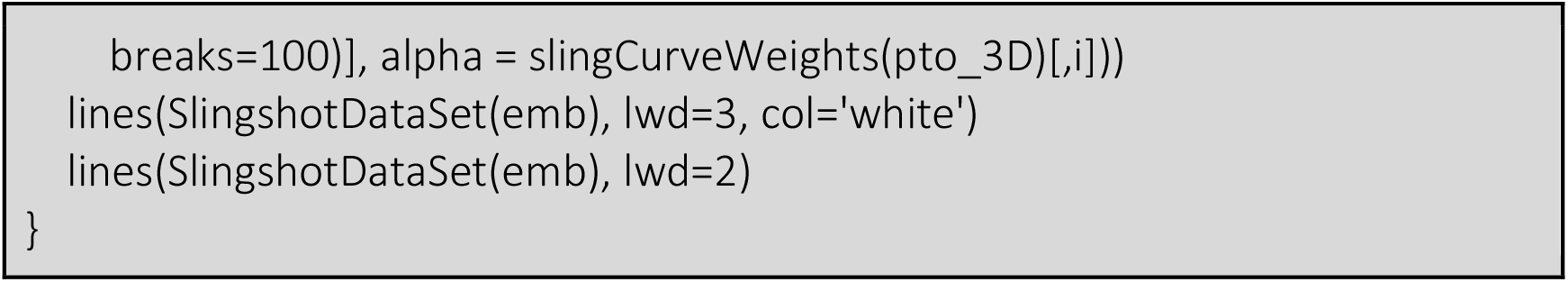

### 5.2 RNA velocity analysis pipeline

Here, we combine all RNA velocity aspects discussed into a single pipeline to run RNA velocity analysis on the dataset of endocrine development in the pancreas [19], which has previously been analysed in [22] as well. For this, we will use *scVelo* [22], the Python package implementation accompanying the dynamical model which also includes an efficient implementation of the steady state model. For documentation and instructions on how to install *scVelo*, see *scvelo*.*org*.

The dataset to be analysed can easily be loaded using *scVelo*:

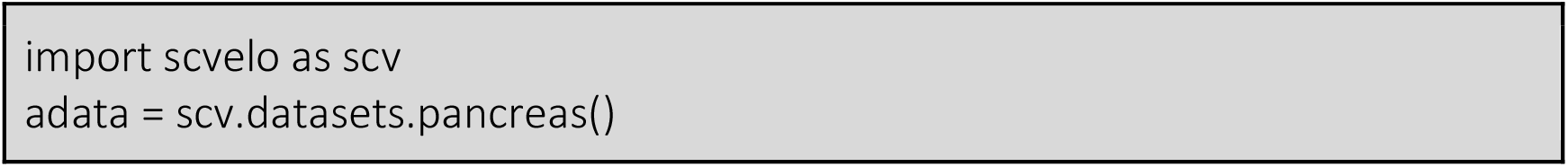

In the resulting AnnData object, estimated unspliced and spliced counts are stored as layers. The X slot, based on which we will perform principal component analysis and calculate the kNN graph, contains the spliced counts as well.

Then, we pre-process the data, as discussed before. Here, we filter genes which share less than 20 counts across modalities, and normalize gene counts by their median. Finally, we only retain the 2,000 most highly variable genes. To mitigate the effect of outliers for PCA and kNN graph calculation, spliced counts are log1p transformed as well, i.e., log(*s* + *1*). The transformation is, however, only applied to the X slot, not the entry in layers. In *scVelo*, we jointly perform all these steps by simply calling the function *pp*.*filter_and_normalize*.

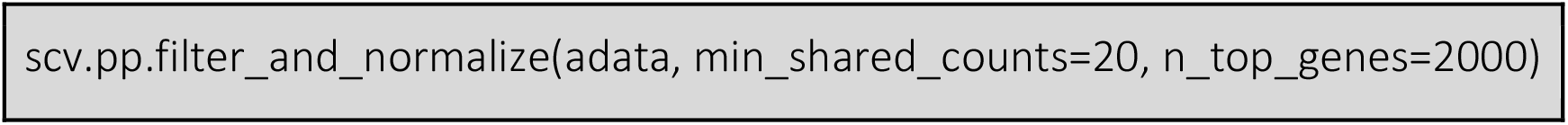

Next, counts need to be imputed by first order moments to reduce the effect of noise. As outlined in Section 3.1, this requires a dimension reduction into PCA space and kNN graph construction. Both steps, including the moment calculation, can be applied using the *pp*.*moments* function.

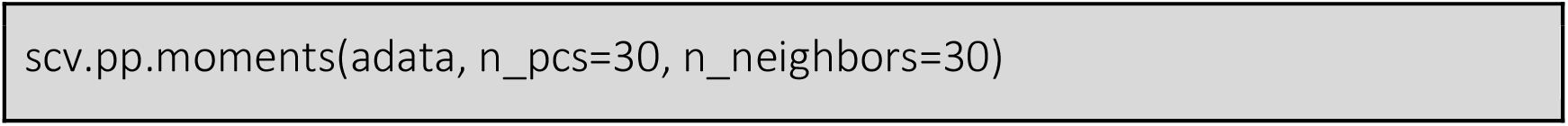

The data is now in a format such that we can estimate RNA velocity. We first calculate velocities using the steady state model, and then with the dynamical one. For the former, we simply need to call the velocity function with *mode=‘deterministic’*.

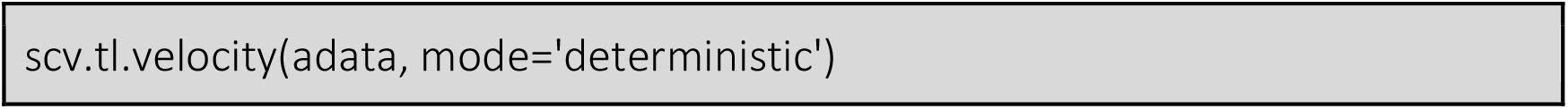

For the dynamical model, we first infer the parameters using *tl*.*recover_dynamics*, then calculate the velocities by specifying *mode=‘dynamical’*. To speed up parameter inference, users can also increase the number of cores used to parallelize calculations, via *n_jobs* parameter; below, we use 4 cores. It is worth noting that setting *n_jobs* to −1 would use all available cores in the machine.

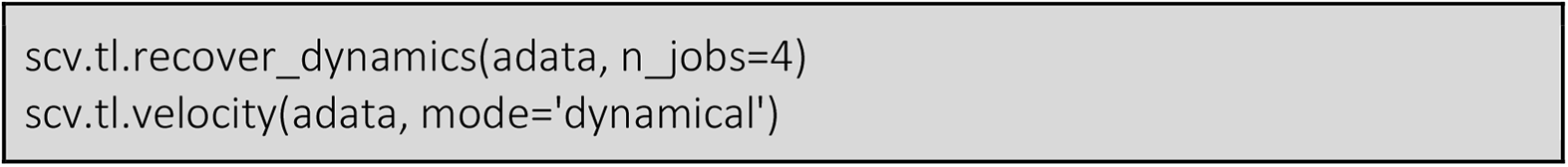

Then, we compute a velocity graph and project the velocity stream onto the UMAP embedding. Note that we did not calculate the UMAP embedding as it is already provided in the AnnData object loaded via *scVelo*. If users were to calculate a UMAP embedding for their dataset, they could, for instance, use the *pl*.*umap* function from the Python package *Scanpy*.

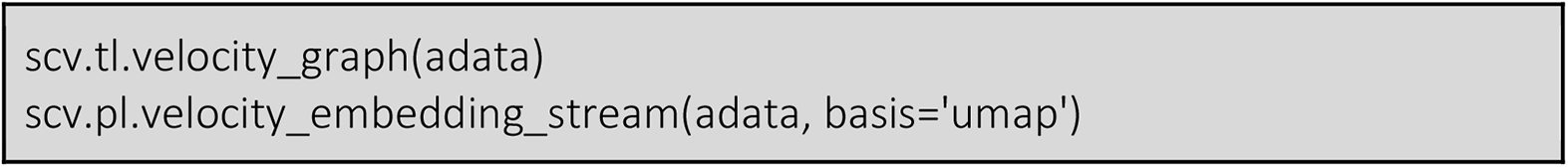

Now, we can compare the two approaches based on their velocity streams. As shown in Figures 10a and 10b, overall, both approaches yield similar results that agree with known cellular differentiation. However, there are also differences between the two cases that are worth mentioning. In the results from the steady state model, there is no flow going into the cluster of Epsilon cells and there is an inconsistent backflow in Alpha cells.

**Figure 10.**
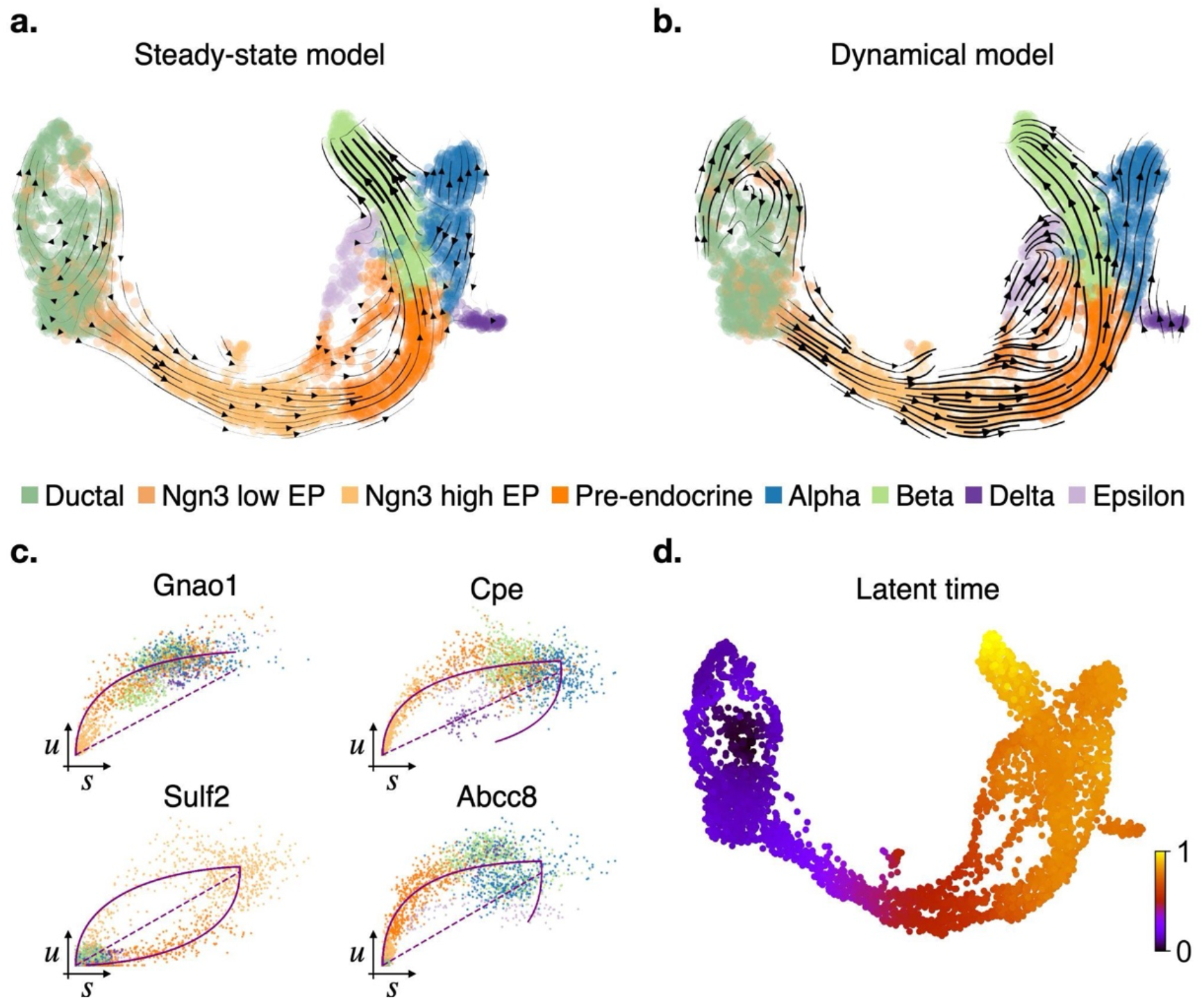
Velocity stream in UMAP-based embedding. **a**. The steady state model does not recover a flow in Epsilon cells and includes an inconsistent backflow in Alpha cells. **b**. Using the dynamical model, flows are more consistent and the dynamics in Epsilon and Delta cells are recovered. **c**. Each gene is assigned a likelihood score indicating its importance in the overall dynamics. Phase portraits of exemplary genes with high likelihood include the inferred trajectory (solid purple line) as well as an estimated steady state ratio (dashed purple). **d**. Gene specific time assignments can be pooled into a single gene-shared latent time, which recovers cellular differentiation in time and agrees with prior biological knowledge.

It is worth highlighting that even in the absence of dynamics, velocity stream embeddings can, and will, display directionality. As such, instead of studying the embeddings, we should focus on phase portraits. One important advantage of the dynamical model (over the steady-state model) is that it provides a likelihood score for each model. Genes with high likelihood contribute more to the overall dynamics compared to those with low scores. In our case, genes with the highest likelihood are informative, i.e., most of their phase portraits display the discussed almond shape curvature. These phase portraits can be plotted using *scVelo*’s *pl*.*scatter* function. Four exemplary genes with high likelihood are shown in Figure 10c. This confirms that RNA velocity analysis can successfully be applied on this dataset.

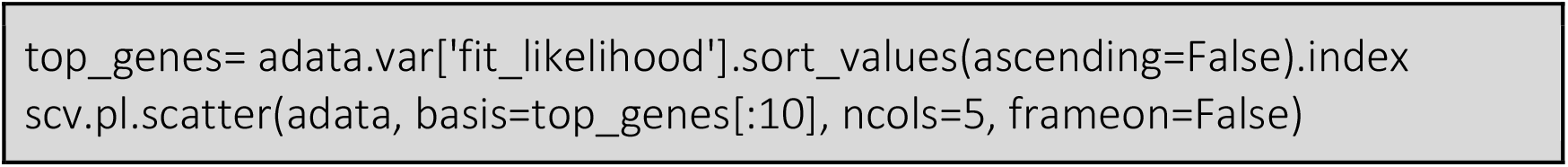

Finally, latent time can be calculated using *scVelo*’s *latent_time* function. As displayed in Figure 10d, the estimated latent time agrees with both prior biological knowledge on cellular differentiation in pancreas, and the velocity stream embedding. Early cell types (Ductal) are assigned early time points, where mature cells (Beta, Alpha, Delta, Epsilon) are associated with later ones.

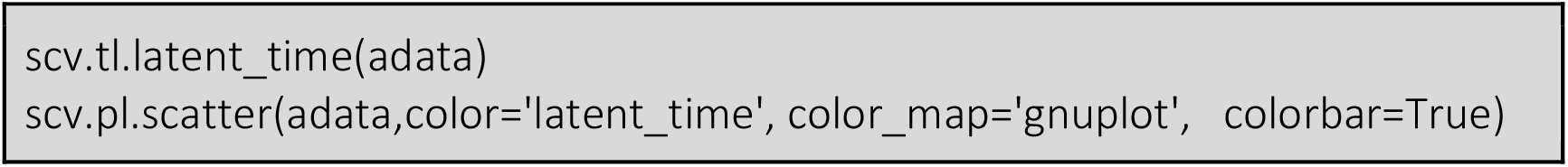

## 6. Notes

1. While we have introduced several types of trajectories, it may not always be clear how to choose among them. For example, the structure of a complex dataset may be overfitted, leading to many bifurcations and/or cycles, while the same dataset could also be described by a more parsimonious trajectory. The balance to strike may be considered analogous to the classical bias-variance trade-off in statistics.
2. Many methods are not flexible enough to allow for different types of trajectories within the same dataset. For example, the pancreas dataset that we have analyzed (Section 4) may be better represented by a combination of a cyclic and a diverging trajectory, however, *Slingshot* is not able to fit such a trajectory.
3. Dynamic processes are subject to change depending on environmental or experimentally induced conditions; the comparison of trajectories between such conditions has recently gained increasing interest. One may compare global features, such as a trajectory’s topology across conditions, as well as smaller changes, such as gene expression across the dynamic, between conditions. It may similarly be of interest to compare RNA velocity estimates across conditions. Such comparisons may hint at, for example, treatment-induced faster progression.
4. To infer RNA velocity, three models exist: steady-state, dynamical and stochastic. The first two have been discussed here, the latter is a generalization of the first (for more details see [22]).
5. The RNA velocity steady-state model assumes splicing dynamics to have reached it equilibria within the experimental time scale, as well as a constant splicing rate across all genes. As a consequence, only the steady-state ratio can be estimated. In contrast to this, the dynamical model solves the full splicing dynamics by considering all available data.
6. For RNA velocity analysis, unspliced and spliced reads need to be pre-processed similarly to standard scRNA-seq workflows (i.e., filtering lowly abundant genes, normalizing counts and subsetting highly variable genes). Furthermore, missing data may be imputed to mitigate the effect of noise.
7. While velocity streams on low dimensional embeddings are useful visualization means, they should not be over interpreted. Instead, phase portraits should be studied first to ensure they exhibit (at least partially) the required almond shape such that RNA velocity can be inferred confidently.
8. To infer RNA velocity using any of the proposed models, the Python package *scVelo* can be used. Instead, *CellRank* should be used for further downstream analyses, such as estimating transition probabilities, automatically identifying initial and terminal states, quantifying fate probabilities as well as gene expression and putative driver genes in different lineages.

## Acknowledgments

The Authors would like to acknowledge Marius Lange and Fabian Theis for their precious comments and suggestions. KVdB is a postdoctoral fellow of the Belgian American Educational Foundation (BAEF) and is supported by the Research Foundation Flanders (FWO) grants 1246220N, G062219N and V411821N.

## Notes

### Competing Interest Statement

The authors have declared no competing interest.

### Summary of Updates

Error in 1st sentence of Section 3.2 fixed.

## References

1. Trapnell C, Cacchiarelli D, Grimsby J, Pokharel P, Li S, Morse M, et al. The dynamics and regulators of cell fate decisions are revealed by pseudotemporal ordering of single cells. Nat Biotechnol. 2014;32: 381–386.

2. La Manno G, Soldatov R, Zeisel A, Braun E, Hochgerner H, Petukhov V, et al. RNA velocity of single cells. Nature. 2018;560: 494–498.

3. Zeisel A, Köstler WJ, Molotski N, Tsai JM, Krauthgamer R, Jacob-Hirsch J, et al. Coupled pre-mRNA and mRNA dynamics unveil operational strategies underlying transcriptional responses to1 stimuli. Molecular Systems Biology. 2011. p. 529.

4. Vallejos CA, Risso D, Scialdone A, Dudoit S, Marioni JC. Normalizing single-cell RNA sequencing data: challenges and opportunities. Nat Methods. 2017;14: 565–571.

5. Kiselev VY, Andrews TS, Hemberg M. Challenges in unsupervised clustering of single-cell RNA-seq data. Nat Rev Genet. 2019;20: 273–282.

6. McInnes L, Healy J, Melville J. UMAP: Uniform Manifold Approximation and Projection for Dimension Reduction. arXiv. 2018.

7. van der Maaten L, Hinton G. Visualizing Data using t-SNE. J Mach Learn Res. 2008;9: 2579–2605.

8. Hicks SC, Townes FW, Teng M, Irizarry RA. Missing data and technical variability in single-cell RNA-sequencing experiments. Biostatistics. 2018;19: 562–578.

9. Townes FW, Hicks SC, Aryee MJ, Irizarry RA. Feature selection and dimension reduction for single-cell RNA-Seq based on a multinomial model. Genome Biol. 2019;20: 295.

10. Srivastava A, Malik L, Smith T, Sudbery Patro R. Alevin efficiently estimates accurate gene abundances from dscRNA-seq data. Genome Biol. 2019;20: 65.

11. He D, Zakeri M, Sarkar H, Soneson C, Srivastava A, Patro R. Alevin-fry unlocks rapid, accurate, and memory-frugal quantification of single-cell RNA-seq data. bioRxiv. 2021.

12. Melsted P, Booeshaghi AS, Liu L, Gao F, Lu L, Min KHJ, et al. Modular, efficient and constant-memory single-cell RNA-seq preprocessing. Nat Biotechnol. 2021;39: 813–818.

13. Luecken MD, Theis FJ. Current best practices in single-cell RNA-seq analysis: a tutorial. Mol Syst Biol. 2019;15: e8746.

14. Diaconis P, Goel S, Holmes S. Horseshoes in multidimensional scaling and local kernel methods. The Annals of Applied Statistics. 2008;2: 777–807.

15. Saelens W, Cannoodt R, Todorov H, Saeys Y. A comparison of single-cell trajectory inference methods. Nat Biotechnol. 2019;37: 547–554.

16. Ji Z, Ji H. TSCAN: Pseudo-time reconstruction and evaluation in single-cell RNA-seq analysis. Nucleic Acids Res. 2016;44: e117.

17. Street K, Risso D, Fletcher RB, Das D, Ngai J, Yosef N, et al. Slingshot: cell lineage and pseudotime inference for single-cell transcriptomics. BMC Genomics. 2018;19: 477.

18. Cao J, Spielmann M, Qiu X, Huang X, Ibrahim DM, Hill AJ, et al. The single-cell transcriptional landscape of mammalian organogenesis. Nature. 2019;566: 496–502.

19. Bastidas-Ponce A, Tritschler S, Dony L, Scheibner K, Tarquis-Medina M, Salinno C, et al. Comprehensive single cell mRNA profiling reveals a detailed roadmap for pancreatic endocrinogenesis. Development. 2019;146.

20. Wolf FA, Hamey FK, Plass M, Solana J, Dahlin JS, Göttgens B, et al. PAGA: graph abstraction reconciles clustering with trajectory inference through a topology preserving map of single cells. Genome Biol. 2019;20: 59.

21. Amezquita RA, Lun ATL, Becht E, Carey VJ, Carpp LN, Geistlinger L, et al. Orchestrating single-cell analysis with Bioconductor. Nat Methods. 2020;17: 137–145.

22. Bergen V, Lange M, Peidli S, Wolf FA, Theis FJ. Generalizing RNA velocity to transient cell states through dynamical modeling. Nat Biotechnol. 2020;38: 1408–1414.

23. Hastie T, Tibshirani R, Friedman J. The Elements of Statistical Learning: Data Mining, Inference, and Prediction. Springer Science & Business Media; 2013.

24. Chari T, Banerjee J, Pachter L. The Specious Art of Single-Cell Genomics. bioRxiv. 2021.

25. Lange M, Bergen V, Klein M, Setty M, Reuter B, Bakhti M, et al. CellRank for directed single-cell fate mapping. bioRxiv. 2020.

26. Haghverdi L, Büttner M, Wolf FA, Buettner F, Theis FJ. Diffusion pseudotime robustly reconstructs lineage branching. Nat Methods. 2016;13: 845–848.

27. Setty M, Kiseliovas V, Levine J, Gayoso A, Mazutis L, Pe’er D. Characterization of cell fate probabilities in single-cell data with Palantir. Nat Biotechnol. 2019;37: 451–460.

28. Barile M, Imaz-Rosshandler I, Inzani I, Ghazanfar S, Nichols J, Marioni JC, et al. Coordinated changes in gene expression kinetics underlie both mouse and human erythroid maturation. Genome Biol. 2021;22: 1–22.

29. Bergen V, Soldatov RA, Kharchenko PV, Theis FJ. RNA velocity-current challenges and future perspectives. Mol Syst Biol. 2021;17: e10282.

30. Zheng GXY, Terry JM, Belgrader P, Ryvkin P, Bent ZW, Wilson R, et al. Massively parallel digital transcriptional profiling of single cells. Nat Commun. 2017;8: 14049.

31. Zappia L LA. zellkonverter: Conversion Between scRNA-seq Objects. R package version 1.4.0. 2021.

